# Rapid changes in Atlantic grey seal milk from birth to desertion

**DOI:** 10.1101/170514

**Authors:** Amanda D. Lowe, Sami Bawazeer, David G. Watson, Suzanne McGill, Richard J.S. Burchmore, P.P (Paddy) Pomeroy, Malcolm W. Kennedy

## Abstract

True seals have the shortest lactation periods of any group of placental mammal. Most are capital breeders that undergo short, intense lactations, during which they fast while transferring substantial proportions of their body reserves to their pups, which they then abruptly desert. Milk was collected from Atlantic grey seals (*Halichoerus grypus*) periodically from birth until near desertion. Milk protein profiles matured within 24 hours or less, indicating the most rapid transition from colostrum to mature phase lactation yet observed. There was an unexpected persistence of immunoglobulin G almost until weaning, potentially indicating prolonged trans-intestinal transfer of IgG. Among components of innate immune protection were found fucosyllactose and siallylactose that are thought to impede colonisation by pathogens and encourage an appropriate gut microbiome. These oligosaccharides decreased from early lactation to almost undetectable levels by weaning. Taurine levels were initially high, then fell, possibly indicative of taurine dependency in seals, and progressive depletion of maternal reserves. Metabolites that could signal changes in the mother’s metabolism of fats, such as nicotinamide and derivatives, rose from virtual absence, and acetylcarnitines fell. It is therefore possible that indicators of maternal metabolic strain exist that signal the imminence of desertion.

## Introduction

Milk is the sole source of nutrition and passive immune protection for neonatal mammals. Milk changes dramatically in composition in the immediate postpartum period from colostrum to mature phase milk that, in eutherians (‘placental mammals’), then changes little until weaning ([1, 2]). That initial transition may take about 48 hours (as in cattle, sheep, camel; [3, 4]), or it can extend to 30-40 days (as in at least one species of bear; [5]). The composition of colostrum varies among species, particularly in the concentration of immunoglobulins (antibodies) that are a sample of those in circulation in the mother. The class of immunoglobulin that predominates in colostrum is a function of the type of placenta possessed by a given species ([2]).

Immunoglobulins are not the only form of maternally-derived immune protection. Others include several anti-microbial proteins and oligosaccharides. The latter may not be digested for energy provision but instead act against colonisation by potentially pathogenic microorganisms by competitively blocking their mucous and cell surface attachment receptors [6, 7]. Importantly, oligosaccharides are also important for the establishment of a gut microbiome appropriate for the neonates of a species (to both aid digestion of milk and compete with incoming pathogens), and can be heterogeneous and polymorphic between individuals [7-12]. Like the proteins present during the colostrum to mature milk transition, oligosaccharides may change in composition with time after birth, some appearing early, then disappearing, and others may show the inverse [5, 13]. The diversity and changes in oligosaccharide content during lactation has, however, been investigated in only a few species.

We recently reported on the dramatic changes in the proteins, oligosaccharides, metabolites and lipids in the species of eutherian mammal with the longest colostrum to mature milk transition known, the giant panda [5, 8, 14]. This prolonged transition time may be associated with the altriciality of panda neonates, which is the most extreme known amongst eutherians, though not as pronounced as in marsupials [15, 16].

We now report on the opposite extreme, in true seals (Phocidae), which give birth to large, precocious pups that are, in many species, nursed without the mother leaving to feed [17]. The pups are typically deserted after a very short lactation, such that weaning is sudden and there is no period of mixed feeding. As a whole, the true seals are remarkable in these highly abbreviated lactation periods relative to their body masses, the most extreme case being hooded seals that lactate for the shortest time known for any mammal, three to five days [17], the longest amongst marine seals being between five and seven weeks in Weddell seals [17]. The lactation strategies of marine phocids are distinct from other pinnipeds, the otariids (sea lions and fur or eared seals) and odobenids (walrus) despite the fact that they occupy superficially similar marine environments and ecological positions (see summary Figure S1). Otariids lactate for considerably longer (4 to 18 months) during which time some mothers cease lactation for periods while foraging in distant feeding grounds, and, remarkably, re-start lactation on their return [15, 17, 18]. Odobenids may nurse for up to two years, and, unusually amongst pinnipeds, nurse their young while at sea [17, 18].

True seals are considered to be capital breeders, in that maternal body reserves are transferred to their neonates with little or no replenishment until weaning [18-20]. During this period of fasting there is a dramatic loss of maternal body mass to fund a doubling of pup body mass [19, 21]. The adaptive advantage of this intense, abbreviated lactation is under debate but represents a strategy by which a capital breeder can rapidly transfer food with reduced energy expenditure associated with foraging [18].

Here we chose a species of true seal with a lactation period before desertion that is in the mid range amongst phocids, and in which females do not forage at sea during lactation. This is the Atlantic grey seal, *Halichoerus grypus*, that lactate for approximately 16 days, though this varies regionally [17, 22], and our population lactate for between 17 and 23 days. In this we had two aims. First, to establish the time course of colostrum to mature phase lactation in a true seal, and, secondly, to seek components indicative of changes in maternal metabolism and potential signals of approaching desertion. We found that the colostrum to milk transition is extremely rapid in this species, in terms of establishment of mature protein and oligosaccharide profiles. On the other hand, we found that other micronutrients and metabolites change more gradually through lactation, some of which may be indicative of alterations in maternal metabolism leading to desertion.

## Materials and Methods

### Milk collection, storage and processing

The seal milk samples were collected from the Isle of May, Scotland, colony of Atlantic grey seals during October and November 2013, and stored frozen until processed. A further collection was made in November 2016 in an attempt to obtain samples as close after birth as possible without risking adverse maternal behaviour or survival of pups; the collection times would have fallen between 10 to 19 hours. Females were tranquilised with ^®^Zolatil, followed by intravenous oxytocin to stimulate milk let-down, and finally an intramuscular prophylactic dose of tetracycline. No deaths or premature desertions of pups followed any samplings. Samples were centrifuged at 3,000 rpm at 4°C in a Heraeus 1.0R centrifuge with swing-out buckets for 15 minutes and the layer between the upper fat layer and the pellet was taken for analysis (see Figures S2 and S3).

### Protein electrophoresis

One-dimensional (1-D) vertical sodium dodecyl sulphate polyacrylamide gel electrophoresis (SDS-PAGE) was carried out using the Invitrogen (Thermo Scientific, Paisley, UK) NuPAGE system with precast 4-12% gradient acrylamide gels, and β-mercaptoethanol (25 μl added to 1 ml sample buffer) as reducing agent when required. Gels were stained for protein using colloidal Coomassie Blue (InstantBlue; Expedion, Harston, UK) and images of gels were recorded using a Kodak imager. Electronic images were modified only for adjustment of contrast and brightness. Pre-stained molecular mass/relative mobility (M_r_) standard proteins were obtained from New England Biolabs, Ipswich, MA, USA.

### Proteomics

Stained protein bands or spots were excised from preparative 1-D or 2-D gels stained with Coomassie Blue and analysed by liquid chromatography-mass spectrometry (LC-MS) as previously described [5]. Protein identifications were assigned using the MASCOT search engine to interrogate protein and gene sequences in the NCBI databases and linked resources. Searches were restricted to the Caniformiae, and allowed a mass tolerance of 0.4 Da for both single and tandem mass spectrometry analyses. BLAST searches, or searches of genome databases within or beyond the Carnivora, were carried out to check the annotations.

### Metabolomics

Ammonium carbonate, HPLC grade acetonitrile, and methanol were purchased from Sigma-Aldrich, UK. HPLC grade water was produced by a Direct-Q 3 Ultrapure Water System from Millipore, UK. The mixtures of metabolite authentic standards were prepared from standards obtained from Sigma Aldrich, UK. In order to analyse the more polar fraction of the milk samples (0.5 mL) were thawed at room temperature and then centrifuged or 10 minutes at 15,000 rpm at 4ºC (Eppendorf 5424 R, maximum RCF = 21.130g). An aliquot of the supernatant (200µl) was mixed with acetonitrile (800µl). The solution was mixed thoroughly, emulsion was centrifuged for 10 minutes at 15,000 rpm at 4ºC (Eppendorf 5424 R, maximum RCF = 21.130g), and the supernatant was transferred to an HPLC vial for Liquid Chromatography-Mass Spectrometry (LC-MS) analysis. The lipids in the milk were analysed by mixing 200 µl of the whole milk with 800 µl of isopropanol. The solution was mixed thoroughly and emulsion centrifuged for 10 minutes at 15,000 rpm at 4°C (Eppendorf 5424 R, maximum RCF = 21.130g). The supernatant was transferred to an HPLC vial for Liquid Chromatography-Mass Spectrometry (LC-MS) analysis.

HILIC–HRMS and multiple tandem HRMS analysis and data processing was carried out on an Accela 600 HPLC system combined with an Exactive (Orbitrap) mass spectrometer (Thermo Fisher Scientific, UK). An aliquot of each sample solution (10μL) was injected onto a ZIC-pHILIC column (150 × 4.6mm, 5µm; HiChrom, Reading UK) with mobile phase A: 20mM ammonium carbonate in HPLC grade water (pH 9.2), and B: HPLC grade acetonitrile. The gradient programme was as follows: 80% B (0 min) 20 % B (30 min) 5% B (36 min) 80% B (37 min) 80% B (45 min). Peak extraction and alignment were calculated by integration of the area under the curve, using MZMine 2.14 software, as previously described [10]. Resulting data were searched against an in house metabolite database. Similar procedures were used for the lipids analysis which was carried out on an ACE Silica gel column (150 x 3 mm, 3 µm particle size) with mobile phase A 20 mM ammonium formate in water isopropanol (80:20) and mobile phase B acetonitrile/isopropanol (20:80). The flow rate was 0.3 mL/min and gradient was as follows: 0–1 min 8% B, 5 min 9% B, 10 min 20% B, 16 min 25% B, 23 min 35% B, 26–40 min 8% B.

## Results and Discussion

### Proteins

We first compared the protein profiles of milk samples taken at intervals postpartum from several seals, and typical results from single mothers are shown in Figures 1 and S4. These show that the mature, main phase lactation pattern appeared very rapidly after birth, with some major protein bands changing in intensity. Establishing the precise times of birth is difficult in the field, but in a subsequent season we were able to obtain samples from mothers that gave birth between 10 and 19 hours before, and compared the protein profiles with those of two 7-day postpartum samples (Figure 2). This emphasised the very rapid development of the mature protein profile, which was essentially complete within a day. The protein bands numbered in Figure 1 were excised from that gel and submitted for proteomics, the results of which are given in Table 1, along with the putative functions of each protein. The identities of the proteins found were provided with additional confidence from a 2-dimensional protein electrophoresis gel (Figure S5 and Table S1).

**Figure 1.**
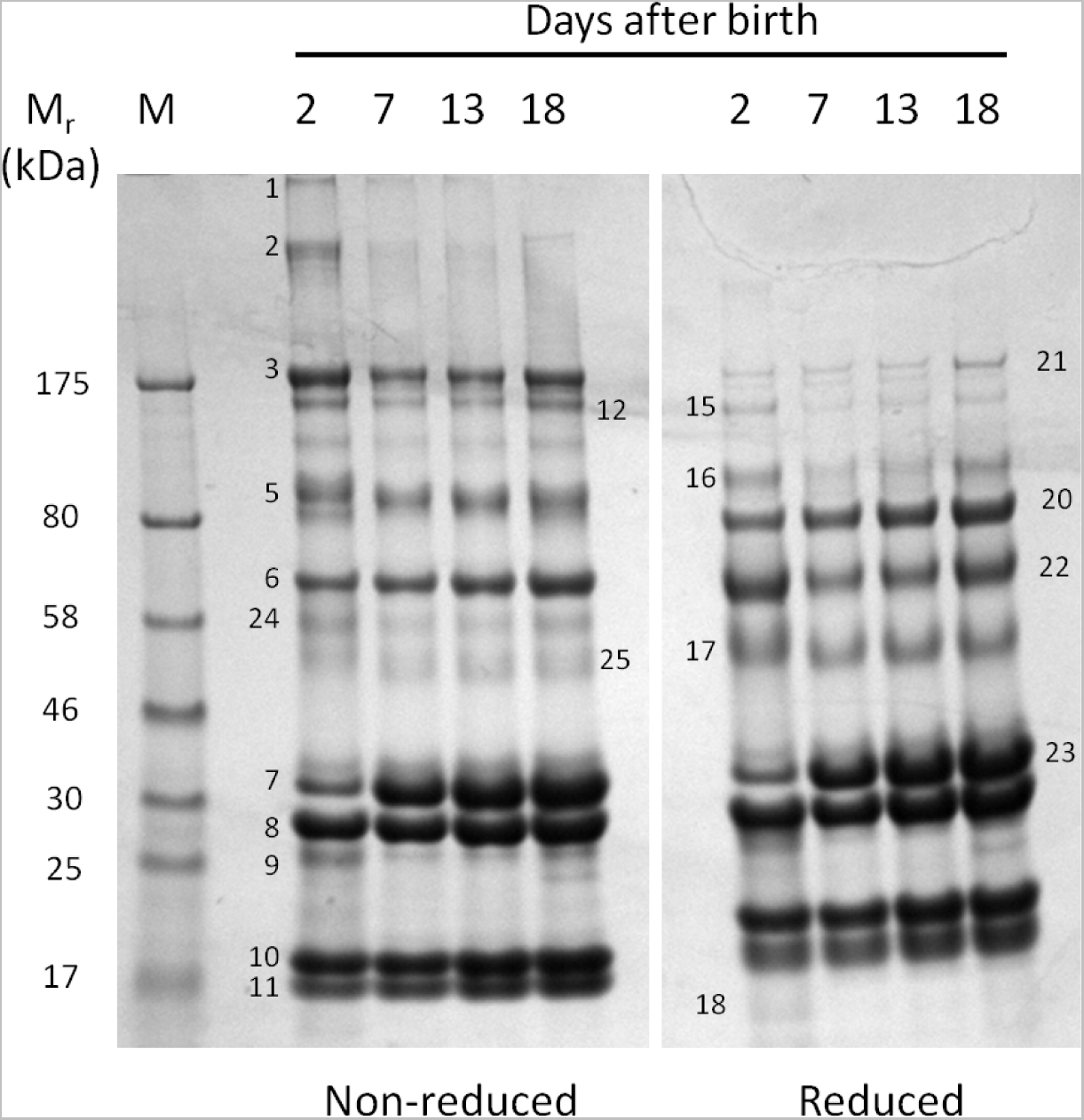
Changing protein profiles of grey seal milk with time after birth. Gradient SDS-PAGE of milk samples obtained from a single mother seal on the days indicated, stained with Coomassie Blue. The protein bands indicated by numbers were excised from the gel and subjected to proteomic identification, the results are given in Table 1. See Figure S1 for a similar protein gel analysis of a sample series from a different seal mother that shows closely similar profiles. M, size reference proteins with their molecular masses given in kiloDaltons (kDa). Samples reduced with β-mercaptoethanol where indicated.

**Figure 2.**
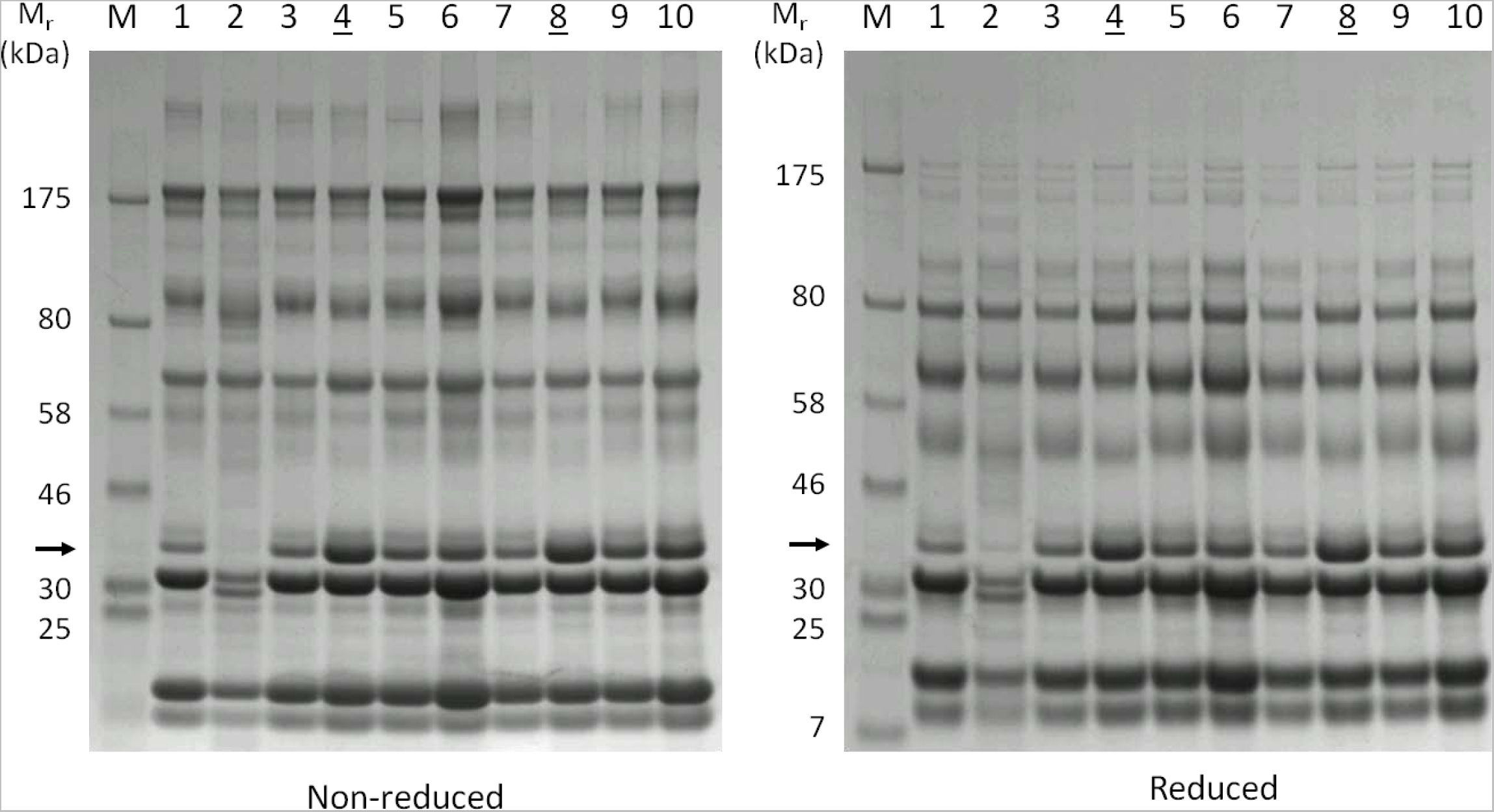
Protein profiles of grey seal milk soon after birth. Milk samples were collected between 10 and 19 hours after birth (numbered tracks), except for tracks 4 and 8 (underlined) which were instead loaded with comparator samples taken 7 days after birth from different mothers in a previous year. Note the absence of the band indicated by the arrow in track 2 and that this band was of lesser intensity in all tracks relative to that in the day 7 samples. The sample in track 2 (and also, to a lesser extent, track 7) also had the smallest fat layer following centrifugation at 4°C (Figure S3). Samples were reduced with β-2-mercaptoethanol where indicated. Different mothers sampled on the Isle of May during November 2016, with those of tracks 4 and 8 taken in November 2014. M, size reference proteins with their molecular masses given in kiloDaltons (kDa).

**Table 1.** Identification of the proteins isolated from bands excised from the protein electrophoresis gel shown in Figure 1.

| Band<sup>a</sup> | Protein<sup>b</sup> | MASCOT score<sup>c</sup> | Number of peptides (unique peptide matches)<sup>f</sup> | Function, association, synonyms and comments<sup>g</sup> |
| --- | --- | --- | --- | --- |
| 1, 16 | Immunoglobulin $\mu$ heavy chain | 108 | 9 (2) | IgM. Antibody. Pentameric. Abundant in serum and colostrum, less so in secretions. |
| 1, 8, 9, 12, 16 | Immunoglobulin $\lambda$ | 116 | 11 (3) | Light chain isoform associated with all immunoglobulin subclasses. |
| 2 | Immunoglobulin $\alpha$ heavy chain | 61 | 11 (1) | IgA. Antibody. Dimeric. Abundant in secretions and milks. |
| 2, 16 | Polymeric immunoglobulin receptor | 457 | 34 (16) | Receptor for IgA and IgM mediating secretion, part of which (secretory component) remains bound to IgA to protect it against proteolytic cleavage in intestine. |
| 3, 12, 15, 22 | Immunoglobulin $\gamma$ heavy chain | 137 | 11 (4) | IgG. Antibody. Most abundant immunoglobulin class in plasma, much less so in secretions. Transferred across placenta or gut in some species by an IgG-specific receptor – situation not known in phocids. |
| 2, 15, 21 | Thrombospondin | 353 | 28 (14) | Extracellular matrix protein. Binds heparin. |
| 5 | Granulins | 174 | 13 (5) | Possible cytokine-like activity. They may play a role in inflammation, wound repair, and tissue remodeling. |
| 6, 20 | Serum albumin | 857 | 108 (39) | Most abundant protein in blood plasma. Carries fatty acids, hydrophobic steroid hormones, hemin, small positively-charged molecules and drugs. |
| 7, 8, 23 | β-casein | 168 | 30 (6) | Phosphoprotein. Source of amino acids, delivers calcium, phosphate, lipids. Structural component and determines the surface properties of the casein micelles. |
| 9, 17, 25 | Apolipoprotein A | 68 | 4 (2) | In plasma, transporter of cholesterol from tissues to the liver and cofactor for the lecithin cholesterol acyltransferase. |
| 10 | β-lactoglobulin-1 | 148 | 16 (5) | Binds and probably transports retinol (vitamin A), vitamin D, and fatty acids including polyunsaturated fatty acids. |
| 11 | Fatty acid-binding protein, heart isoform | 113 | 12 (4) | Thought to play a role in the intracellular transport of long-chain fatty acids and their acyl-CoA esters. Syn. mammary-derived growth inhibitor. |
| 12, 22, 24, 25 | Lactadherin | 580 | 42 (18) | Maintains intestinal epithelial homeostasis and the promotion of mucosal healing. |
| 12, 15 | Ceruloplasmin | 126 | 9 (4) | The major copper-carrying protein in the blood, and plays a role in iron metabolism. Possibly involved in pulmonary antioxidant defence. |
| 17 | κ-casein | | | Stabilizes milk micelle formation, prevents casein precipitation. |
| 17 | α-1-acid glycoprotein<br>(syn. orosomucoid) | 66 | 5 (2) | Acute phase protein in blood. Levels change in pregnancy and in acute phase (fever) response. Binds negatively-charged small molecules, steroids, proteinase inhibitors. Immune regulation. |
| 18 | α-lactalbumin | 47 | 1 (1) | Regulatory subunit of lactose synthase. Changes the substrate specificity of galactosyltransferase making glucose a good acceptor substrate for this enzyme enabling lactose synthase to synthesize lactose. |
| 22 | Xanthine dehydrogenase | 496 | 49 (20) | Key enzyme in purine degradation. Contributes to the generation of reactive oxygen species. Involved in milk fat globule secretion and also innate immunity. |
| 25 | Haptoglobin | 69 | 8 (2) | Indicator of infection or inflammation. Acute phase protein. Captures free haemoglobin. Anti-microbial. |

<sup>a</sup> Gel band codes as indicated in Figure 1.
<sup>b</sup> Protein identifications. Peptides matching to keratin, were excluded.
<sup>c</sup> MASCOT (MOWSE) search score where scores greater than 38 are taken to be significant. The MASCOT score given is the highest value obtained where the protein was identified in more than one band, as were the peptide match values.
<sup>f</sup> Number of peptides found to match with the number of peptides unique to this identification in parentheses.
<sup>g</sup> Putative functions and comments are drawn from literature cited, or NCBI and UniProtKB/Swiss-Prot databases.
All proteomics data and protein diagnostics has been deposited in the Dryad Digital Repository. At the time of writing there are limited genomic, mRNA and protein sequence data available for the grey seal and the protein identifications in this table derive mainly from searching within the Caniformia, the best fits arising from these species - *Leptonychotes weddelli* (Weddell seal), *Pusa hispida* (ringed seal), *Odobenus rosmarus divergens* (walrus), *Ailuropoda melanoleuca* (giant panda), *Mustela putorius furo* (ferret/European polecat), *Ursus maritimus* (polar bear), *Canis familiaris* (domestic dog), *Neovison vison* (American mink), *Arctocephalus tropicalis* (subantarctic fur seal). The database accession codes for the best fits are also given in the Dryad Digital Repository files.

The main proteins segregate between those for adaptive and innate immune protection, and those for nutritional support. Among the former were three immunoglobulin classes (IgG, IgM and IgA), as detected by the presence of their eponymous heavy chains, along with their associated light chains. The immunoglobulins generally appeared in greatest amounts early after birth, such as seen in Figure 1. They were accompanied by the polymeric immunoglobulin receptor that mediates the trans-epithelial transport of immunoglobulins into secretions, predominately IgA, which it then protects against proteolytic cleavage [23-25]. In all species, IgA appears to be continuously present in both colostrum and mature phase milk, presumably to protect the mammary gland and the oral and gastrointestinal tracts of the neonate [25, 26]. IgA tends to be the predominant immunoglobulin in the colostrum of species in which trans-placental transfer of IgG occurs (such as in humans and rodents, which have haemochorial placentae [27, 28]) using the FcRn transporter system [29, 30]. In contrast, IgG tends to be particularly enriched in the colostrum of species in which trans-placental transfer does not occur (e.g. cattle, sheep, horses, camels; epitheliochorial placentae; [27, 28]) [29-31]. In these species, IgG (along with IgM) crosses the gut epithelia directly into the neonate’s circulation for the short period before the gut cell layer closes (24 hours post-partum or less), it then appears at much lower levels from the time at which the colostrum period ends (~24-36 hours) [31].

The zonary discoid endotheliocorial placentae of many Carnivora have peripheral haemophagous zones through which transfer of large plasma proteins such as immunoglobulins may occur, possibly by pinocytosis and phagocytosis of maternal blood rather than mediated by FcRn [32, 33]. Among the Carnivora, trans-placental transfer of IgG occurs to a limited degree in dogs [34], but apparently not in cats [35], but we know of no published information on this for pinnipeds. There is therefore no certainty that passive antibody immunisation occurs before birth in true seals; analysis of the serum of neonates before suckling would, of course, be definitive. Surprisingly, IgG appears to persist at high levels throughout lactation in grey seal milk (Figures 1, 2 and S4). In some mammals, such as rats [29], IgG is actively transported across the gut mucosa (using the same FcRn receptor system as for trans-placental transfer [29]), so it may be that this also applies to seals. If so, then this would be an unusual adaptation in seals that might relate to immune protection of a rapidly growing pup that will soon be deserted and exposed to infections circulating in a breeding colony.

Several proteins of innate immunity were detected. Xanthine dehydrogenase/oxidase is found in most mammal milks and is thought to be defensive, but it also has a role in lipid synthesis and secretion [36, 37]. α-1-acid glycoprotein, ceruloplasmin and haptoglobin were also found and are among a set of proteins that rapidly appear in greatly enhanced amounts in blood at the onset of an acute phase (fever) reaction in mammals [38, 39]. They are usually synthesised in the liver, but it is now known that some acute phase proteins can be synthesised in mammary gland tissue in response to infections, and then appear in milk [40-42]. An inflammatory response in mammary gland tissue is observable during phases of the lactation cycle when the gland is undergoing reconstruction and may be in a vulnerable state [43]. The presence of protective proteins in grey seal milk could therefore be due to microbes colonising an active mammary gland, or as a prophylactic against infection.

The other main proteins found are well-established as being specialised for milk-based nutrition, such as the caseins [44-46]. β-casein was present at lower levels in the earliest samples relative to day 7 (Figures 1, 2 and S4), and even missing in one (arrowed in Figure 2; Figure 1, band 7). A delayed post-parturient appearance of caseins has also been observed in the giant panda, in which secretion of both β- and κ-caseins may take 30-40 days to reach main phase levels [5]. β-casein is a highly phosphorylated protein that transports calcium ions and forms micelles that appear to be stabilised by κ-casein [47]. The delayed appearance of caseins may explain how the soluble, fat-depleted layer of early grey seal milk samples is less turbid (milky) than later ones (Figure S2), as is also the case in giant pandas [5].

β-lactoglobulin was present at high relative levels in all samples, including those collected soon after birth (Figures 1 and 2; Table 1). It is present in all Carnivoran milks that have been examined, in which it may occur in one to three isoforms [5, 48]. It is thought to be a carrier of long chain fatty acids and retinol (Vitamin A) [48, 49]. Retinol is insoluble and highly sensitive to oxidation but can be protected within an apolar protein binding site [48, 50-53] (and M.W. Kennedy, unpublished). Retinoic acid derivatives of retinol are crucial to a wide range of cell differentiation and developmental processes in vertebrate [54-60], so the safe delivery of its precursor to a rapidly growing neonatal seal may be particularly important. Curiously, humans (and camels, elephants) [5, 61] do not produce β-lactoglobulin, though some primates do (macaques and baboons) [48], so its true role in milk remains mysterious.

Proteinase inhibitors were also found. A specific colostrum trypsin inhibitor is present in many mammal milks, the concentration of which appears to correlate positively with that of IgG [62]. In bovine milk, for example, this inhibitor is found for only 2 – 48 hours postpartum, which fits with the idea that it is there to reduce cleavage of immunoglobulins undergoing transfer to the neonate. The encoding gene has been examined in otariids and odobenids in which it appears to be functional, but it is disrupted in one phocid (Weddell seal) [62]. If this is also true in the grey seal, then its absence in our survey is explicable, but this then begs the question of whether the other proteinase inhibitors we found act to compensate for protection of the unusually prolonged secretion of IgG into the milk of this species.

Two proteins that are more usually associated with blood plasma were present, albumin and apolipoprotein A, both of which are involved in lipid transport in blood, albumin carrying a range of small charged molecules in addition to fatty acids. Whether these two proteins are made in, or actively transported from blood by, the mammary gland, or leak passively into milk from blood plasma, remains to be established, though the high level of albumin present suggests an influence of some kind in milk. A general, non-specific leakage of blood plasma components into the milk is unlikely given that we did not find other major plasma proteins such as complement C3 or transferrin.

α-lactalbumin was found, which is interesting given its role in lactose production (see below).

### Oligosaccharides

Complex sugars are abundant in the milk of many species, though not all, and are active as free or protein-linked oligosaccharides [12, 63-66]. In humans, these complex sugars vary dramatically in quantity and types between mothers [12, 67]. They are generally not digested to provide a neonate’s energy metabolism but are instead thought to control colonisation by pathogens through interfering with their sugar-based adhesion mechanisms required for binding to mucus layers or cell surfaces [7, 68]. Milk oligosaccharides also play a crucial role in establishing an appropriate microbiome by, for instance, acting as a selective nutrient supply for species of *Bifidobacterium* [7, 69, 70].

We found that both fucosyllactose and sialyllactose were present soon after parturition in grey seal milk but were then rapidly lost with time after birth, until little or none of either was detectable towards the end of lactation (Figure 3). Sialyllactose (N-acetylneuraminyllactose) occurs in 3’ and 6’ forms, the former being the most common in milks. Our MS analysis indicated that only one form was present in the seal milk, and that was probably 3’. The differences in amounts of these sugars varied considerably between mothers in the first week, which could indicate intrinsic differences between the mothers in how much they produce, or the rates at which secretion of these oligosaccharides change with time after birth. Levels of these two complex sugars decreased roughly simultaneously, which is the opposite to the trend found in the giant panda [5]. In that species, fucosyllactose rose with time, but the 3’ form of sialyllactose fell. The rate of change in the concentrations of these oligosaccharides in seal milk was very much greater in seals than in giant pandas, in which it takes 20 to 60 days at least for levels of these oligosaccharides to stabilise [5].

**Figure 3.**
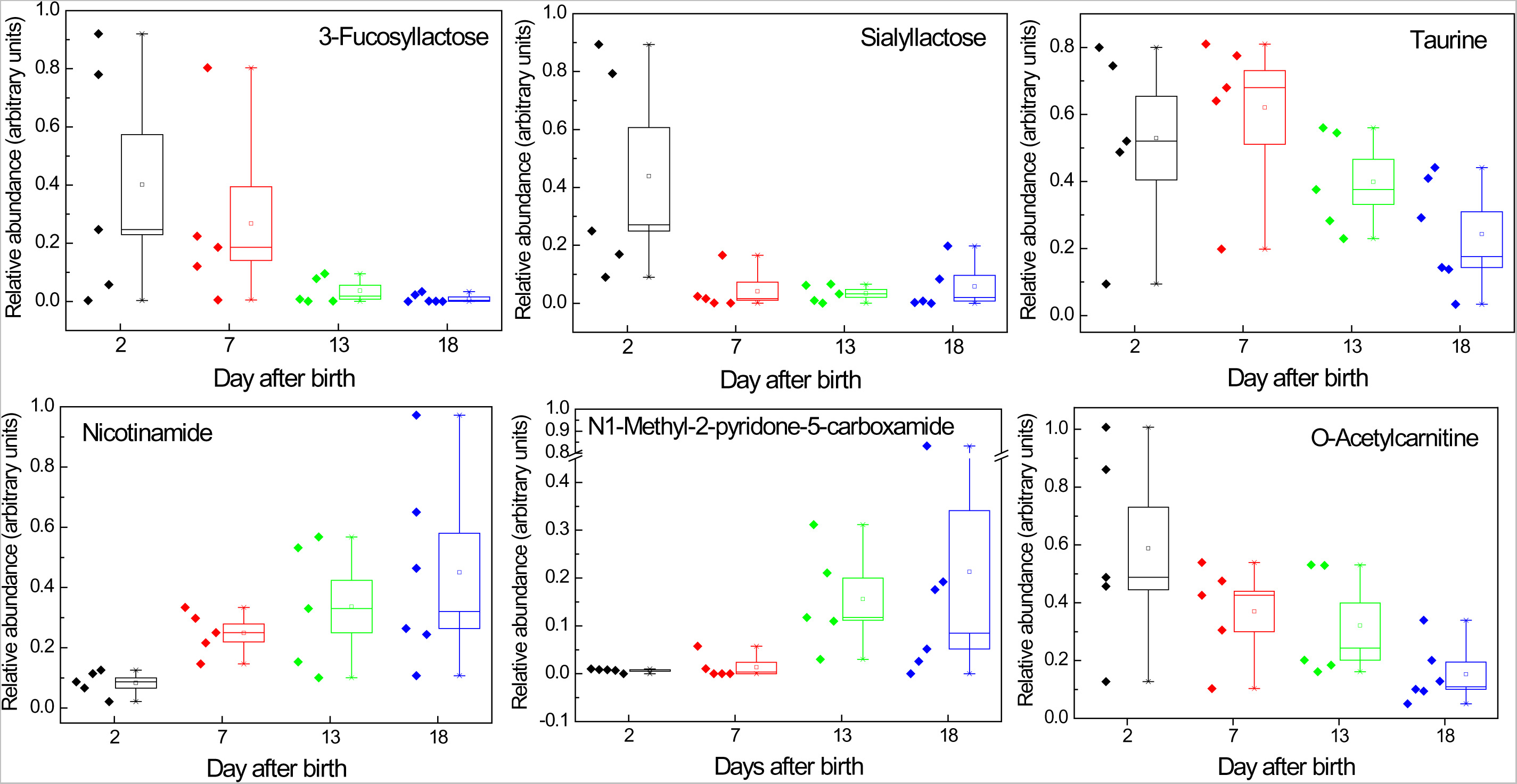
Changes in grey seal milk metabolites and micronutrients with time after birth. Selection of compounds exhibiting changing concentrations as lactation progresses. Fucosyllactose and sialyllactose are oligosaccharides that control colonisatio by microbes. Taurine is an essential dietary requirement in some species of hyperpredator. The remaining metabolites are associated with fat metabolism, potentially pertinent to nursing seals in terms of mobilisation of their body reserves, and lipids required for rapid growth and maintenance of pups that need to accumulate large fat reserves for the forthcoming starvation period, and for subsequent thermal insulation at sea. The data indicated for 18 days after birth are a mixture from samples taken on days 17, 18 and 19. The square symbol in a box is the mean; the band in the box is the median; the box extends to the standard error of the mean; the whiskers indicate the range.

Lactose is the principal energy component of the milk of many species of land mammal (e.g. cow, sheep, horse, dog, camel, human), but is either at very low levels or absent in marine mammals [2, 18]. Lactose is found at very low levels in phocids, but is absent in the milks of otariids and odobenids [2, 18]. This loss is postulated to be because lactose’s role in energy provision is completely supplanted by milk fats, and that one of lactose’s functions, the osmotic drawing in of water into milk [71, 72], is not advantageous in marine mammals [73]. Lactose is synthesised by lactose synthase, which is a two-component enzyme comprising β1,4-galactosyltransferase (which is produced in many tissues) and α-lactalbumin (which is specific to mammary glands). Otariids and odobenids appear to have alterations to their α-lactalbumin–encoding gene that would disable the protein’s enhancement of lactose synthesis - which is not the case in phocids [73]. Despite finding α-lactalbumin in grey seal milk (see Figure 1 and Table 1), lactose was present but only in amounts that are very low relative to those in cow, goat and camels (data not shown), consistent with studies on other phocids [71, 73]. In true seals, therefore, the lactose may instead be there to provide a substrate for the synthesis of the fucosylated and sialated forms of lactose that are for management of the gut or mammary gland microbiome, or protection against microbial pathogens, rather than for energy supply [74].

### Taurine

Taurine has a multitude of biological functions, such as involvement in membrane stabilisation and modulation of calcium signalling, and it is essential for cardiovascular function, development and function of skeletal muscle, the retina, and the central nervous system [75-77]. In addition there is increasing evidence that taurine is essential for supporting the immune system since it is found at very high levels in phagocytes [78]. Moreover, human neonates have a limited capacity to produce taurine [78]. One of the primary bile acids of mammals is taurine-conjugated, so a rich supply of it may be crucial for the processing of a fat-rich diet, which particularly applies to the neonates of marine mammals. In that regard, bile salts also activate bile salt-activated lipase that is involved in digestion of lipids [79], and is found in grey seal milk (Table S2). Some species of hypercarnivore, such as cats and possibly also polar bears [80-82], cannot synthesise taurine, and are thereby dependent on dietary sources. As we will report elsewhere, we find that taurine occurs at considerably higher concentrations in seal milk than in milks of many other species. Being piscivorous hypercarnivores that have ready access to plentiful sources of taurine in their diet, seals, like other hypercarnivores, may have foregone synthesising taurine, which would then be an essential requirement in their milks. Here, we found that the concentration of taurine is, like other small molecules, highly variable in milk samples from mother to mother, but is highest soon after birth and then falls as weaning approaches (Figure 3). If grey seals cannot synthesise and replenish taurine, then that reduction could be due to depletion in the mother during her fast, which should not apply to those phocids in which the females periodically forage during lactation [17].

### Micronutrients or indicators of metabolic activity?

We examined changes in metabolites that are involved directly in, or are indicative of, fat-fuelled energy metabolism, and have here selected nicotinamide, acetylcarnitine and N1-methyl-2-pyridone-5-carboxamide for note. As we will report elsewhere, we find that nicotinamide, its derivatives and precursors (such as anthranilic acid) are dramatically higher in concentration in seal milk than in a selection of land mammals (cow, goat, camel), that this also applies to N1-Methyl-2-pyridone-5-carboxamide, and some carnitines.

Nicotinamide is required for the production of NAD^+^, which is a key co-factor in fatty acid β-oxidation. Since the energy metabolism of both seal mothers and pups is based on large scale oxidation of fats, then a high requirement for NAD^+^ would be expected, and we found that the concentration of nicotinamide increases with time of lactation (Figure 3). As with taurine and oligosaccharides, there is substantial diversity in milk nicotinamide levels between mothers at all four sampling times, which could relate to their initial nutritional states, physiological condition, or demand for milk by their pups. As with other small molecule metabolites, the increasing concentrations of nicotinamide could be a reflection of the need for the pups to be supplied. Or that a mother’s own fat metabolism is increasingly drawn upon, as she continues her fast, and nicotinamide leaks into her milk from her blood circulation.

Nicotinamide can also be converted to N-methylnicotinamide, which has in the past been viewed as a non-biologically active waste product, but is increasingly attracting interest as a stimulator of peroxisome proliferation [83-85], which is pertinent to a fasting mother seal - the metabolism of long chain fatty acids takes place in peroxisomes before transfer to the mitochondria. N-methylnicotinamide is metabolised into N1-methyl-2-pyridone-5-carboxamides via the action of aldehyde oxidase and also cytochrome P450 2E1 (CYP2E1), and it has been proposed that its levels give an indication of peroxisome proliferation [83, 86, 87]. N1-methyl-2-pyridone-5-carboxamide is only present at very low levels at the beginning of lactation and increases dramatically with time until the end of lactation (Figure 3). This compound could therefore be an indicator of increasing fat metabolism in the mothers and possibly a potential marker of when a mother may soon depart that may be detectable in both blood and milk.

Carnitine is centrally involved in fatty acid metabolism and fulfils three main functions - it transports fatty acids into mitochondria so that they can undergo β-oxidation to generate NADH; it removes fatty acids from the mitochondria in order to maintain the levels of free CoA within a certain range; and it removes waste fatty acids from the body as water soluble carnitine conjugates [88]. As we will report elsewhere, carnitines that are conjugated with long acyl chains (e.g., oleoyl, palmitoyl, and docosahexanoyl in particular) are substantially more abundant in seal milk than in cow, goat or camel milks, whereas those conjugated with short acyl chains (acetyl, propionyl, butyl) were of similar abundances or slightly lower. However, the post-parturition changes in seal milk were similar for all types, and Figure 3 illustrates the trend for acetylcarnitine, which diminishes to low levels towards the end of lactation.

As for the other small molecules that we found in seal milk, we cannot be sure whether the carnitines are there to supplement a pup’s metabolic activity or whether they are reflecting a mother’s physiology at the time of sampling, or both. Dietary carnitine is an important contributor to the carnitine pool and short chain acyl forms may have improved bioavailability in comparison to free carnitine. Also, acylcarnitines are activated for metabolism by mitochondria since they can be converted directly to acyl CoA with the investment of a molecule of ATP which is required for the conjugation of free acyl groups to CoA [89, 90]. Long chain fatty acids such as docosahexenoic acid are metabolised in peroxisomes to shorter chain acids before entering the mitochondria for further metabolism. They are required for conversion to acyl CoAs before they can be oxidised in the peroxisomes and again it would be advantageous if they were available in their activated form e.g. docosahexanoyl carnitine. Thus, aside from whether or not the acylcarnitines can be efficiently absorbed by seal pups, for every molecule of acyl carnitine assimilated a molecule of ATP is conserved.

Amongst food sources derived from animals, carnitine is most abundant in red meats, followed by fish and milk. Given the extremely high dependence of seal pups on fats, it is perhaps not surprising that they are provided with such high levels of acylcarnitines, and that maternal provision early in lactation would be valuable. It is interesting, though, that, whilst carnitine levels drop overall with time, other metabolites involved in fatty acid metabolism and long chain acyl carnitines increase (e.g. nicotinamide). This perhaps reflects the use of carnitine in the formation of the “ready to go” acyl carnitines and the requirement for nicotinamide for NAD+ formation to support β-oxidation after their conversion to acyl CoAs.

## Conclusions

There is no widely accepted definition of what colostrum is. We previously defined the point at which colostrum ends and main phase lactation begins as being when the components of milk stabilise in relative concentrations [5]. We find that there is no such point in the brief lactation period of grey seals. We have therefore here taken the end of colostrum as being when the protein profiles have stabilised.

The transition from colostrum to main phase lactation in the Atlantic grey seal is the shortest yet recorded for any species of mammal. It is in stark contrast to the longest known for a eutherian, that which occurs in a fellow member of the Carnivora, a bear [5]. This divergence is all the more impressive given that true seals, along with other pinnipeds, share membership of the Caniformia suborder within the Carnivora [91, 92]. It is conceivable that the transition occurs even more quickly in species of seal in which the lactation period is even shorter, the hooded seal in particular.

Our focus has been on the components involved in immune defence and indicators of metabolic changes. The rapid change in protein profile is particularly impressive, but so too is the persistence of IgG with time after birth. This is unusual and could indicate a particular need to provision a rapidly growing offspring with a sufficient supply of antibody to maintain its defence against pathogens in circulation in breeding colonies, the phocine and other morbilliviruses being obvious examples [93, 94]. A question therefore is whether this prolonged delivery of IgG is only for protection of the gut, or instead results in a systemically protective build-up of this immunoglobulin in the blood of the pups before desertion. Of innate immune protection, the changes in oligosaccharides are also of note. Those probably involved in antimicrobial activity were present only at the beginning of lactation, and many fewer types were found in comparison to bears. The differences between the composition and changes in milk oligosaccharides between two species within the Carnivora suggest stark differences in their adaptations to pathogen defence and the microbiomes they need to establish.

We observed changes in compounds central to fat metabolism that could either be reflections of how the mother’s metabolism alters as she mobilises and transfers her own body resources to her pup without replenishment, or donation of compounds to aid the pup’s own fat metabolism, or both. Either way, our findings merit optimism in finding a metabolic indicator of when a seal mother reaches the end of her resources and must leave.

## Acknowledgements

We are grateful to Scottish National Heritage for permitting our time and sampling permission on the Isle of May.

## Author contributions

Conceived and developed the project – MWK, PPP

Carried out the analyses – ADL, SB, DGW, SMc, RJSB, MWK

Analysed the data – ADL, SB, DGW, SMc, RJSB, MWK

Wrote the paper – MWK, PPP, DGW, RJSB in consultation with all authors.

## Ethical statement

Collection of milk samples was approved by the ethical committee of Scottish Oceans Institute, and the College of Medical, Veterinary and Life Sciences Ethics Committee of the University of Glasgow.

## Access to data

All the proteomics and metabolomics data will be deposited in the Dryad Digital Repository, until which any requests for data should be sent to the corresponding author.

**Supplementary information accompanies this paper** – see separate file

